# Proving a negative; estimating species ‘Confidence in Absence for Decision-Making’ (CIADM) using environmental DNA monitoring

**DOI:** 10.1101/2024.05.12.593768

**Authors:** Nathan P. Griffiths, Bernd Hänfling, Marco Cattaneo, Rosalind M. Wright, James A. Macarthur, Sara Peixoto, Jonathan D. Bolland

## Abstract

1. Policy-driven decision-making is an important aspect of environmental management globally, often focused on protecting priority species. However, declining trends in freshwater biodiversity have resulted in a lack of up-to-date knowledge regarding the distribution of rare and elusive species. Furthermore, when dealing with priority species, it is sometimes more important to provide a confident assessment of absence, than merely confirm their presence. Without such confident assessments, resource intensive management plans may be misplaced, and not adequately targeted to conserve important remaining populations.
2. Here, we present a framework to estimate confidence in absence, referred to as “Confidence in Absence for Decision-Making” (CIADM), based on single-visit environmental DNA metabarcoding data obtained from water samples. It uses a case study of European eel presence / absence upstream of 44 water pumping stations, given their critically endangered status and the legislative drivers for remediation (EC Eel Regulation 1100/2007, Eels (England and Wales) Regulation 2009). Through a high degree of biological (sample) and technical (PCR) replication, we retrospectively assigned ‘confidence in absence’ values and proposed various strategies to achieve the required confidence levels in future surveys.
3. Our findings indicate that 17 out of 44 pumping stations tested positive for eel, and we were able to assign a >99% confidence level that the remaining 27 sites were negative for eel DNA at the time of sampling. Increasing both biological and technical replication increased ‘confidence in absence’ values. For example, using three PCR replicates per sample, required four replicate biological samples to achieve >95% and six to achieve >99% confidence in eel absence given non-detection. However, we estimate that by using seven PCR replicates per sample a >99% confidence in eel absence following non-detection could be achieved with only three replicate biological samples. Furthermore, we found that eel positive sites had significantly higher species richness, and fish communities differed between eel positive and eel negative sites.
4. This study highlights the importance of optimising workflow specific replication, and provides an adaptable framework to produce confidence estimations of priority species absence given non-detection.

## 1. Introduction

European freshwater ecosystems face widespread challenges such as habitat fragmentation, flow alteration and climate change (Belletti *et al*., 2020; Bolland *et al*., 2019; Szabolcs *et al*., 2022). In response to these multiple interacting pressures, policies have been implemented to aid environmental management and decision-making at various levels. Examples of major initiatives in Europe include the EU Water Framework Directive (WFD; EC, 2000) which is driven by the protection of overall habitat quality (European Environment Agency, 2018), and the EC Eel Regulation (1100/2007) which is driven specifically by priority species. Freshwater biodiversity declines and disparities for conservation across Europe (Szabolcs *et al*., 2022) also underscore the need for up-to-date information on freshwater biota, including the distribution of priority species to meet conservation objectives. However, establishing confidence in the absence of rare and elusive priority species, according to a level of sampling effort, is inherently challenging.

Collecting comprehensive freshwater biodiversity data can be time-consuming and costly (Murdoch *et al*., 2007). As a result, environmental DNA (eDNA) metabarcoding of water samples has emerged as a promising approach to support these efforts (Blancher *et al*., 2022). Substantial work has already been devoted to integrating eDNA-based monitoring into fish-based WFD ecological status assessments in lakes (Willby *et al*., 2019) and rivers (Hering *et al*., 2018; Pont *et al*., 2019). Several studies have found increased sensitivity of eDNA-based monitoring for both entire fish communities (Griffiths *et al*., 2020; McColl-Gausden *et al*., 2021) and specific priority species (Muha *et al*., 2021; Weldon *et al*., 2020) when compared with traditional methods. However, to successfully integrate molecular methods into aquatic monitoring, clear best practice guidelines and an adaptive approach to the implementation of new methods are necessary (Blancher *et al*., 2022).

Environmental DNA metabarcoding based assessment of species presence/absence can be affected by false positives (detection of absent species) and false negatives (failing to detect species that are present) (Buxton *et al*., 2021). Processes to monitor and mitigate false positives are well established, including the use of blanks/negative controls at each stage of the workflow, as well as setting low-frequency reads thresholds (Hänfling *et al*., 2016) and adopting conservative occupancy approaches (Stauffer *et al*., 2021). Increasing replication can help to reduce the rate of false negatives. For example, Ficetola *et al*. (2015) found that optimal levels of PCR replication varied with detection probability, recommending at least eight PCR replicates when the detection probability is expected to be low, and that sample replication from the same locality could further increase detection probability. However, without prior knowledge of a specific workflow, study system, or target species, predicting detection probability and relative importance of biological (sample) and technical (PCR) replication can be challenging. Therefore, such findings may not always be transferable.

Spatiotemporal sampling replication has also been observed to obtain significantly more biodiversity than PCR replication (Beentjes *et al*., 2019). However, in the context of targeted monitoring at priority sites and structures it is important to maximise single-visit point sampling detectability, and single-point sample replication studies are lacking. Elsewhere, multispecies site occupancy models have been successfully applied to spatially replicated eDNA metabarcoding data using binary presence/absence data and more recently variants including sequence reads (Fukaya *et al*., 2022; McColl-Gausden *et al*., 2021). However, occupancy modelling can lead to inaccurate conclusions when detection probability for target species is very low (Ficetola *et al*., 2015). Furthermore, modelling using spatially replicated samples may be inappropriate when the impacts of structures are locally specific (i.e. catchment fragmentation and specific abstraction points).

Overall, there is an urgent need to establish a workflow specific ‘confidence in absence’ framework to effectively prioritise resources, especially in the context of species conservation. One such species is the European eel (*Anguilla anguilla*), a catadromous fish species (Schmidt, 1923) that has experienced significant declines over the last four decades (Aalto *et al*., 2016; Bilotta *et al*., 2011; Correia *et al*., 2018; Jacoby & Gollock, 2014; Podgorniak *et al*., 2016). With *A. anguilla* recruitment now estimated to be a fraction of its 1970s baseline level - 1.4% in the North Sea and 5.6% elsewhere in 2019 (ICES, 2019), the International Union for Conservation of Nature (IUCN) has classified the species as ‘critically endangered’ (Jacoby & Gollock, 2014). Given this alarming decline, the related policy (EC Eel Regulation 1100/2007) includes requirements for eel passage and screening of water intakes, including pumping stations. However, due to the lack of current knowledge about eel distribution, implementing conservation measures in line with policy is challenging. Particularly given that artificial riverine structures are often barriers to species movement, potentially leading to many fragmented and low-density populations. In these circumstances, limited resources must be urgently targeted on the structures which directly impact target species, and not misdirected by prioritising sites where they are absent. For example, in pumped river catchments, downstream passage often requires eels to pass through pumping stations (Bolland *et al*., 2019), which can cause mortality (Buysse *et al*., 2014). Providing safe downstream passage is technically challenging and expensive (Buysse *et al*., 2015), and thus it is important to prioritise remediation efforts (Solomon & Wright, 2012). Environmental DNA metabarcoding has already been validated in pumped catchments, and was particularly sensitive to *A. anguilla* detections when compared with traditional methods (Griffiths *et al*., 2020). However, to avoid erroneously discounting sites for mitigation, and to comply with regulations, the confirmation of a priority species presence is not enough, we must also confidently identify sites where priority species are genuinely absent.

This study therefore develops a framework to assess Confidence in Absence for Decision-Making (CIADM) based on single visit, point sampling eDNA metabarcoding data (Figure 1). The aim is to develop a framework which assigns a confidence of absence, through a combination of biological (sample) and technical (PCR) replication, enabling stakeholders to minimise the risk of false negative surveys. False negatives can be introduced at the biological level (i.e. no DNA collected) or technical level (i.e. no DNA detected). Studies investigating the impact of single point eDNA sample replication are lacking, and so it was important to initially over sample. By doing so, the conditional probability of DNA presence in a sample given occupancy at site (p*_s_*), and the conditional probability of DNA detection in a PCR replicate given presence in sample (p*_r_*), can be estimated. With these, alongside the overall probability of occupancy at site (p*_o_*), CIADM can then discern the impact of biological and technical replication under given circumstances to estimate confidence in absence (Figure 1).

**Figure 1.**
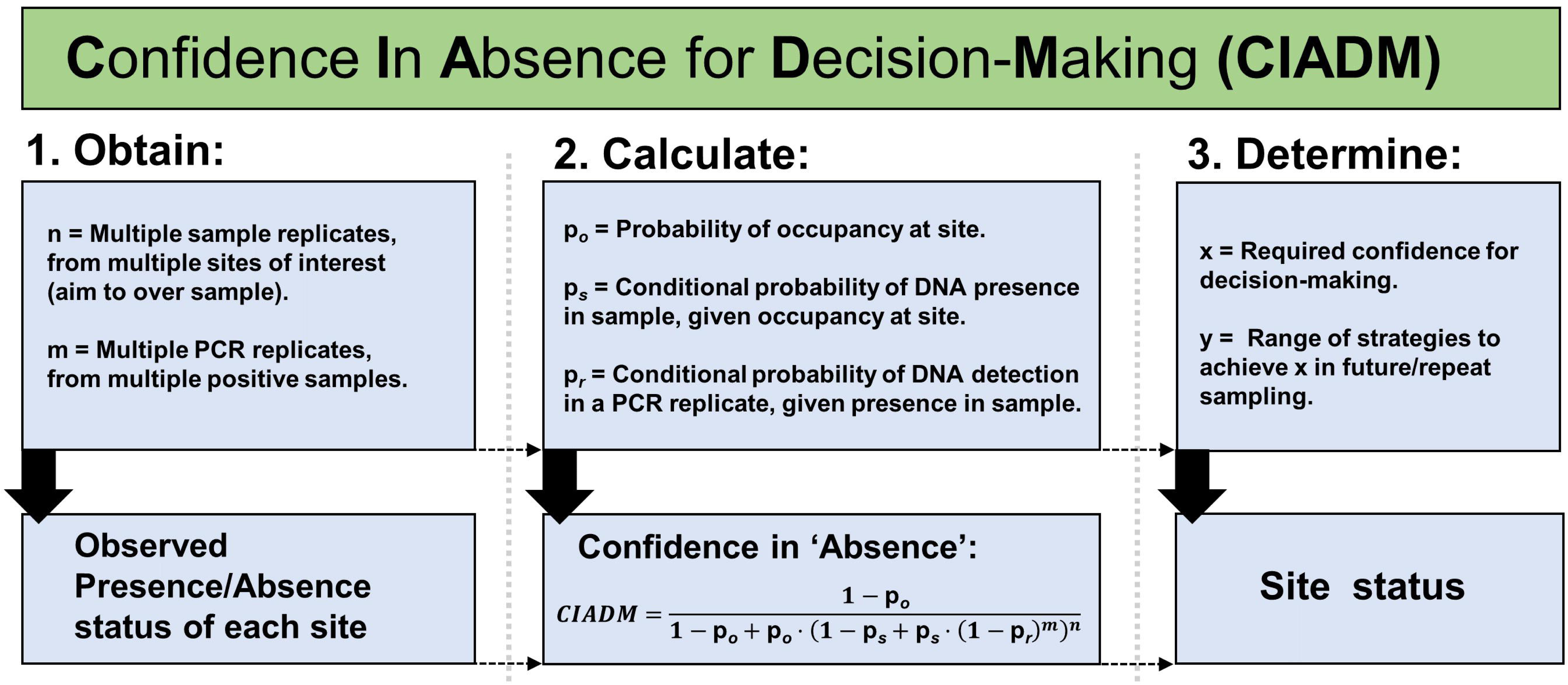
An overview of the confidence in absence for decision-making (CIADM) framework. Including; 1. Data collection and initial classification of sites, 2. Components of the model to calculate confidence in absence and 3. Handling and applications of outputs.

## 2. Methods

### 2.1 Study sites

Water samples were collected from the upstream river catchments of 44 water pumping stations located in Eastern England, UK (Figure 2). These pumping stations were chosen to encompass a range of sizes and geographical characteristics, aiming to provide a representative sample of pumped catchments across the UK. In addition to the standard sampling methodology which involved ten samples taken from each site (total samples = 440), a further PCR replication test was carried out on samples taken from Chain Bridge (CB) (Figure 2), since this site had known presence of our priority species and a 50% initial detection rate with our standard workflow (Section 2.4).

**Figure 2.**
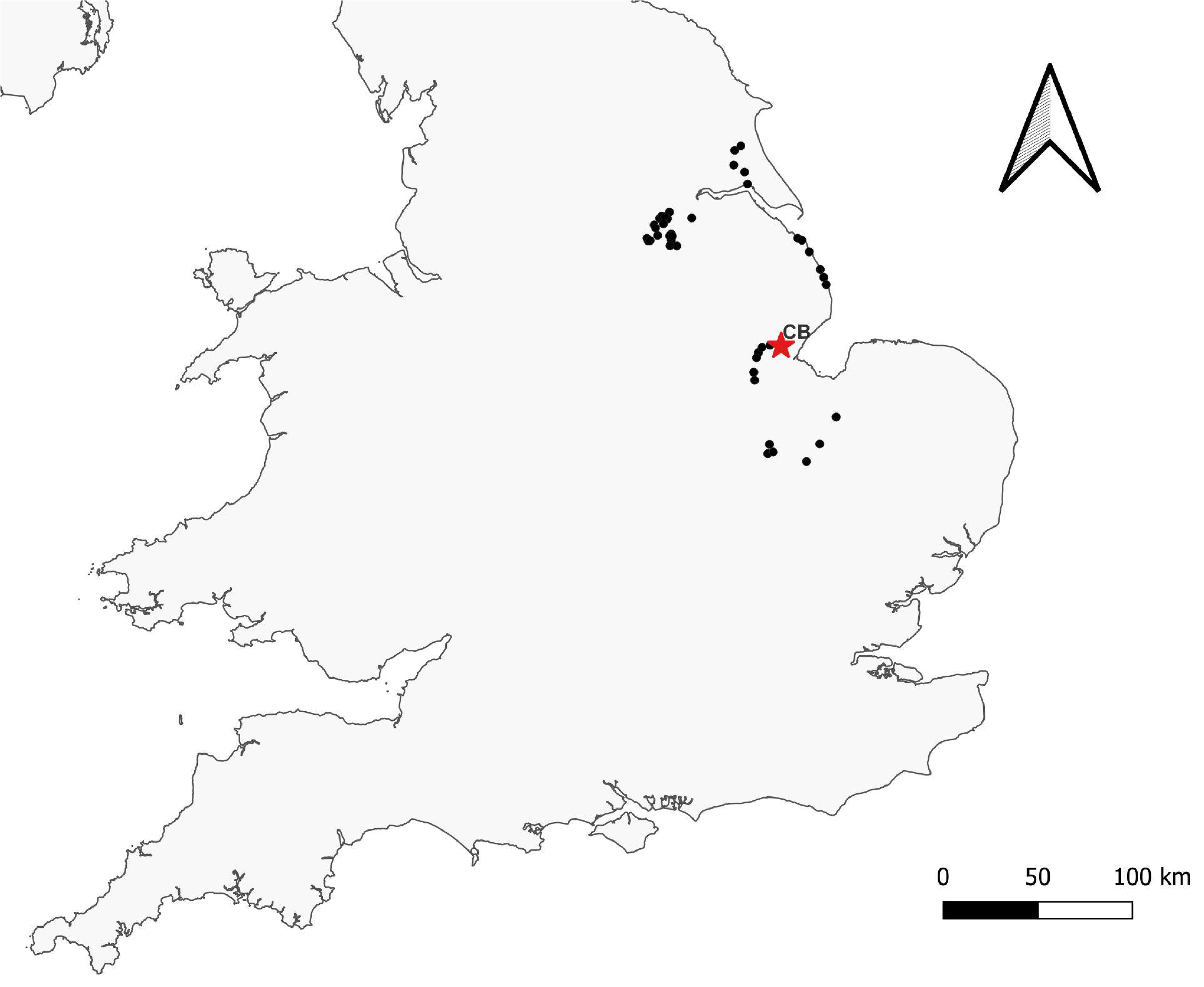
An overview of the sampling region within the UK. Showing the location of each pumping station sampled (black dot), and highlighting the location where PCR replication tests were carried out (red star).

### 2.2 Sample collection

Ten 2L water samples were collected directly upstream of each pumping station structure using sterile Gosselin HDPE plastic bottles (Fisher Scientific UK Ltd., Loughborough, UK). Each sample was taken over a 10m transect, and consisted of 5 x 400ml sub-samples to account for stochastic distributions of eDNA within the watercourse. Sterile gloves were worn by the sampler at each sampling site, and changed between sites. For each site visit, a 2L blank, consisting of a sampling bottle filled with purified water, was taken out and handled alongside eDNA water samples in order to monitor for field contamination. Upon collection, water samples were placed on ice in a bleach-sterilised cool box during transit, and taken back to a dedicated eDNA filtration facility at the University of Hull.

All samples and blanks were vacuum filtered within 24h of collection. Surfaces and equipment in the filtration laboratory were sterilised using 10% v/v chlorine-based commercial bleach solution (Elliott Hygiene Ltd., Hull, UK). Between filtration runs, filtration equipment was immersed in 10% bleach solution for 10 min, soaked in 5% v/v MicroSol detergent (Anachem, Leicester, UK) for an additional 10 min, and following this rinsed thoroughly with purified water to remove any detergent residue. Where possible, the full 2L of each water sample was filtered through sterile 0.45μm cellulose nitrate membrane filters with pads (47 mm diameter; Whatman, GE Healthcare, UK) using Pall filtration units; two filters were used per sample to reduce filter clogging. Once 1L of water had passed (or after 30 minutes under vacuum) the filters were removed from units using sterile tweezers, rolled, and placed back-to-back in sterile 5ml Axygen screw cap transport tubes (Fisher Scientific UK Ltd., Loughborough, UK) each pre-prepared with 1 g of 0.15 mm and 1 - 1.4 mm diameter sterile garnet beads ready for extraction (Sellers *et al*., 2018), then stored at -20°C.

### 2.3 DNA extraction

DNA from each sample was co-extracted from duplicate filters alongside extraction blanks at the University of Hull eDNA facility using a designated sterile extraction area, following the Mu-DNA: Water protocol (Sellers *et al*., 2018). Following extraction, the eluted DNA extracts (100μl) were quantified and checked for purity using a NanoDrop™ Spectrophotometer to confirm that DNA was isolated successfully, then stored at -20°C until PCR amplification.

### 2.4 Library preparation

Library preparation was carried out following the methodology outlined in Griffiths *et al*. (2023), summarised below:

Nested metabarcoding following a two-step PCR protocol was performed, using multiplex identification (MID) tags in both steps to enable sample identification as described in (Kitson *et al*., 2019). The first PCR (PCR1) was performed in triplicate (3× PCR replicates per eDNA extract), amplifying a 106bp fragment using published 12S ribosomal RNA primers 12S-V5-F (5′-ACTGGGATTAGATACCCC-3′) and 12S-V5-R (5′-TAGAACAGGCTCCTCTAG-3′) (Kelly *et al*., 2014; Riaz *et al*., 2011). These primers have been previously validated in silico, in vitro and in situ for UK freshwater fish, confirming that all UK species can be detected with the exceptions of distinctions between *Lampetra planeri* / *Lampetra fluviatilis* and *Perca fluviatilis* / *Sander lucioperca*; three species of Asian carp (*Hypophthalmichthys nobilis*, *Hypophthalmichthys molitrix* and *Ctenopharyngodon idella*); and species within the genera Salvelinus and Coregonus (Hänfling *et al*., 2016). PCR-negative controls (Molecular Grade Water) were used throughout, as were positive controls using DNA (0.05 ng μl^-1^) from the non-native cichlid *Maylandia zebra*. All PCR replicates were pooled, and samples from each PCR1 plate normalised and pooled to create sub-libraries. Sub-libraries were then purified using MagBIND RxnPure Plus magnetic beads (Omega Bio-tek Inc., Norcross, GA, USA), following a double size selection protocol (Quail *et al*., 2009). Ratios of 0.9× and 0.15× magnetic beads to 100μl of amplified DNA from each sub-library were used. Following this, a second shuttle PCR (PCR2) was performed on the cleaned product to bind Illumina adapters to the sub-libraries. A second purification was then carried out on the PCR2 products with Mag-BIND RxnPure Plus magnetic beads (Omega Bio-tek Inc., Norcross, GA, USA). Ratios of 0.7× and 0.15× magnetic beads to 50 μl of each sub-library were used. Eluted DNA was then refrigerated at 4°C until quantification and normalisation. Once pooled, the final library was then purified again (following the same protocol as the second clean-up), quantified by qPCR using the NEBNext Library Quant Kit for Illumina (New England Biolabs Inc., Ipswich, MA, USA) and verified for fragment size and purity using an Agilent 2200 TapeStation with High Sensitivity D1000 ScreenTape (Agilent Technologies, Santa Clara, CA, USA). Once verified, the library was loaded (mixed with 10% PhiX) and sequenced on an Illumina MiSeq using a MiSeq Reagent Kit v3 (600 cycle) (Illumina Inc., San Diego, CA, USA) at the University of Hull.

After sequencing, sub-libraries were demultiplexed to sample level using a custom Python script. Tapirs, a reproducible workflow for the analysis of DNA metabarcoding data (https://github.com/EvoHull/Tapirs), was subsequently used for taxonomic assignment of demultiplexed reads. Sequence reads were quality trimmed, merged, and clustered before taxonomic assignment against a curated UK fish reference database (Hänfling *et al*., 2016). Taxonomic assignment here used a lowest common ancestor (LCA) approach based on basic local alignment search tool (BLAST) matches with minimum identity set at 98%.

For the PCR test site (CB) which presented 5/10 eel positive samples (50%) with the above protocol, each individual sample was re-run with 10x individually sequenced PCR replicates (total replicates over 10 samples = 100). This process followed the same procedure; however, each replicate was individually tagged, demultiplexed, and taxonomically assigned independently in order to investigate the influence of technical (PCR) replication within the described workflow.

### 2.5 Data processing and analysis

Data were analysed and visualised using R version 4.2.1 (R Core Team, 2020). Raw data from the 44 sites were merged into one data frame and CIADM applied based on a subset of 7 species which occurred across standard surveys and PCR replication tests (Section 2.6). In order to apply the model, it is important that biological replication is considered from standard surveys, and also technical replication is considered from PCR tests, in our case those carried out at site (CB). In this study, data for both of these were available for *Anguilla anguilla*, *Cobitis taenia*, *Rutilus rutilus*, *Scardinius erythrophthalmus*, *Esox lucius*, *Gasterosteus aculeatus* and *Pungitius pungitius*.

Following this, the full dataset of standard samples (total=440) was carried forward for further analysis. Prior to any statistical analysis, species richness data were screened for normality using base R. Differences in species richness between priority sites were tested for significance using a nonparametric Wilcoxon rank sum test, while differences in fish community were tested for using PERMANOVA based on Jaccards and Bray-Curtis index. PERMANOVAs were performed using the function ‘adonis’ in the R package vegan (Oksanen, 2013), with 999 permutations. The homogeneity of multivariate dispersions between groups was also calculated using the BETADISPER function, and tested for significance using ANOVA. To enable visualisation, species accumulation curves were plotted for each site using the package vegan (Oksanen, 2013), then box plots and Non-Metric multidimensional scaling (NMDS) using the package ggplot2 (Wickham, 2016).

### 2.6 Confidence in absence framework (CIADM)

We obtained 440 samples (n) over 44 sites and fed these into the CIADM framework (Figure 1). In addition to these, 100 PCR replicates (m) which were individually sequenced from the 10x samples at the PCR test site (CB) were also included, in order to discern the impact of technical replication. Based on the three-level occupancy model approach detailed in (Nichols *et al*., 2008) and (Mordecai *et al*., 2011), the following were estimated:

p*_o_* = probability of occupancy at a site
p*_s_* = conditional probability of DNA presence in a sample given occupancy at the site
p*_r_* = conditional probability of DNA detection in a replicate given presence in the sample

In our study, the conditional probabilities above were considered to be constant for ease of interpretation (Schmidt *et al*., 2013). From these data, the maximum likelihood (ML) estimates of p*_o_*, p*_s_* and p*_r_* were calculated using the expectation-maximization algorithm (Dempster *et al*., 1977). These were then combined to obtain the ML estimate of CIADM: i.e., of the conditional probability of absence given no DNA detection in n samples with m PCR replicates each (Figure 1). The raw data and R code used to apply this model is made available here (See: *Data Availability Statement*).

## 3. Results

### 3.1 Site overviews

Across 44 study sites, the fish species richness ranged from 1 - 15 and the average was 7 (S.D. = 3.9). Seventeen sites (38.6%) tested positive for our priority species, *A. anguilla*. The species accumulation curves visualise how increasing the number of single-point sample replicates provided diminishing returns in regard to detecting new species (Figure 3).

**Figure 3.**
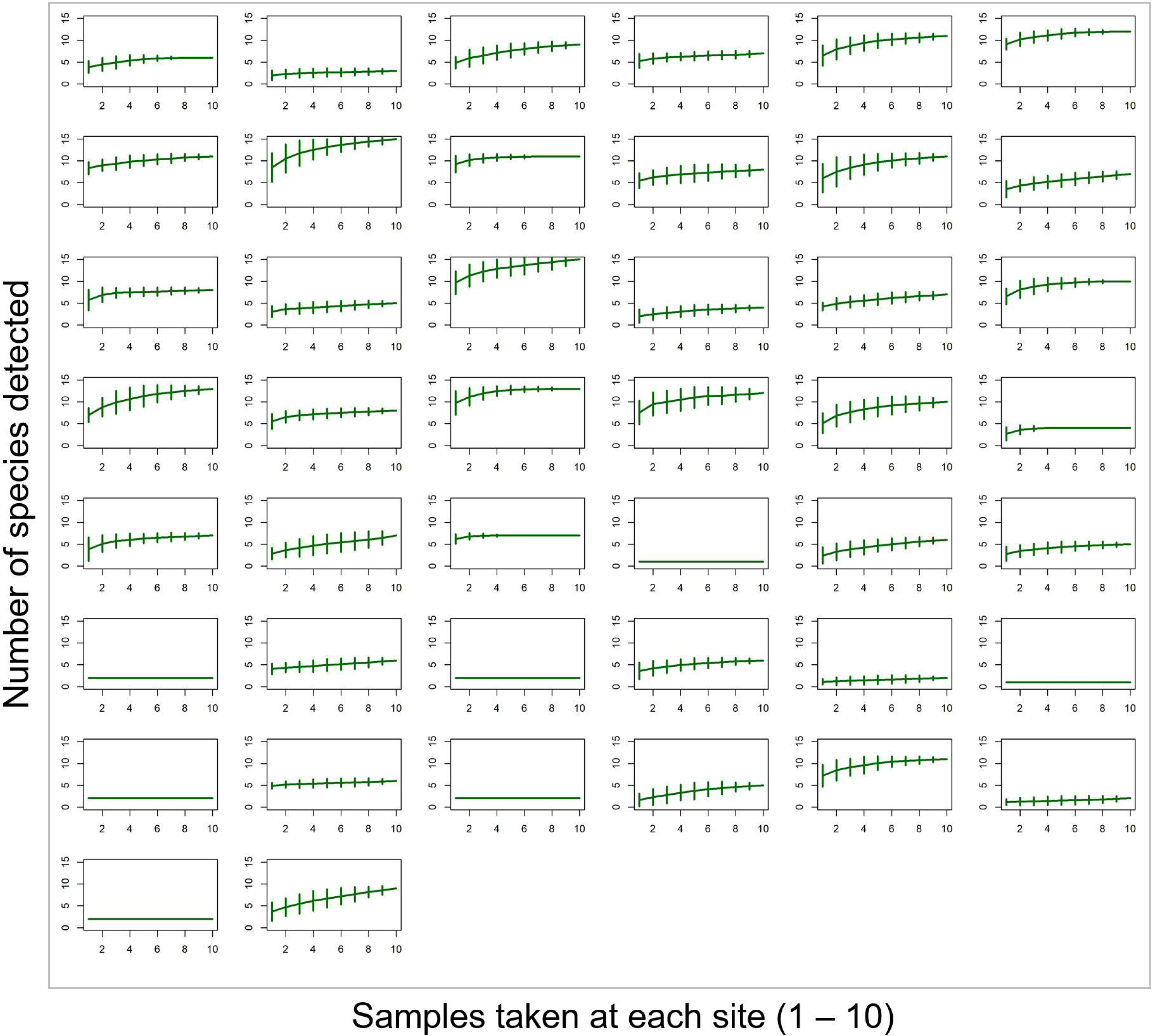
Species accumulation curves (SAC) of the ten samples taken at each of the 44 study sites. Green lines indicate the mean SAC, while vertical whiskers show the standard deviation based on 100 random permutations of the data.

### 3.2 PCR replication tests

At the PCR test site (CB), where 5/10 (50%) of samples initially tested positive for eel using the standard metabarcoding workflow (Section 2.4), eel detection increased to 10/10 (100%) detection when PCR replication was increased to ten replicates per sample. The average number of positive PCR replicates at this site was 3.9 per water sample (SD = 1.66, Range = 2 - 7).

### 3.3 CIADM framework outputs

Based on the seven fish species inputs considered in the CIADM framework, we found that the initial ten samples with three PCR replicates were sufficient to be >99% confident in our assignments of species absence (supplementary information). This confirms that we have over-sampled in our initial survey, meeting the criteria of *n* in CIADM^1^ (Figure 1). In fact, using three PCR replicates, >95% confidence could be achieved with four samples, and >99% with seven samples for all species. In addition, with sufficient PCR replication, only three sample replicates were required to achieve >95% confidence in absence, while four samples were required to attain >99% confidence for all species. The full range of CIADM^2^ outputs for 200 combinations of biological and technical replication for each of our seven study species is available in (supplementary information).

Considering our priority species, *A. anguilla*, the values of p*_o_* = 0.39, p*_s_* = 0.87 and p*_r_* = 0.26 were obtained, producing the range of CIADM^2^ outputs below (Figure 4). Indeed, assuming maximum *n* = 10 and maximum *m* = 20, the CIADM^2^ output provided 139 combinations of replication strategies to achieve >99% confidence in absence given non-detection for *A. anguilla* (Figure 4). In our study, using three PCR replicates per sample, it took four replicate samples to achieve >95% and six to achieve >99% confidence in eel absence given non-detection.

**Figure 4.**
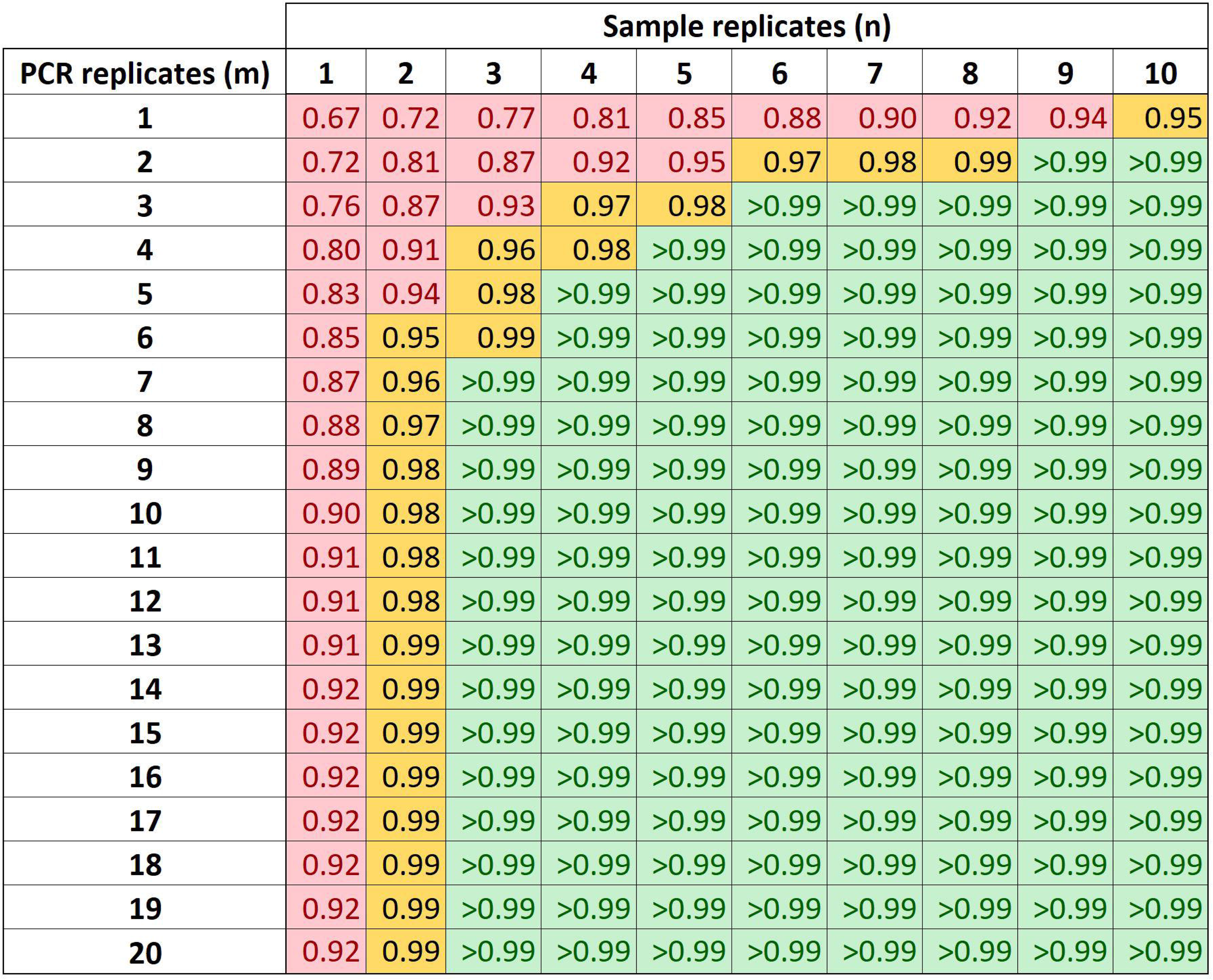
Outputs of the CIADM framework for the European eel, showing the confidence in absence estimations corrected for sampling/lab effort. These outputs were generated from the data obtained at the 44 study sites. Cells are coloured based on confidence values of <95% (Red), 95-99% (Amber) and >99% (Green).

### 3.4 Associated fish community

Based on confirmed site status, we observed that overall species richness was significantly higher at eel positive sites (Wilcoxon rank sum test; W = 35154, p < 0.001) (Figure 5). The overall fish community also varied significantly between sites classified as eel positive and eel negative; Jaccard index (PERMANOVA, R2 = 0.06, DF = 1, P < 0.001) and Bray-Curtis (PERMANOVA, R2 = 0.11, DF = 1, P < 0.001) (Figure 6).

**Figure 5.**
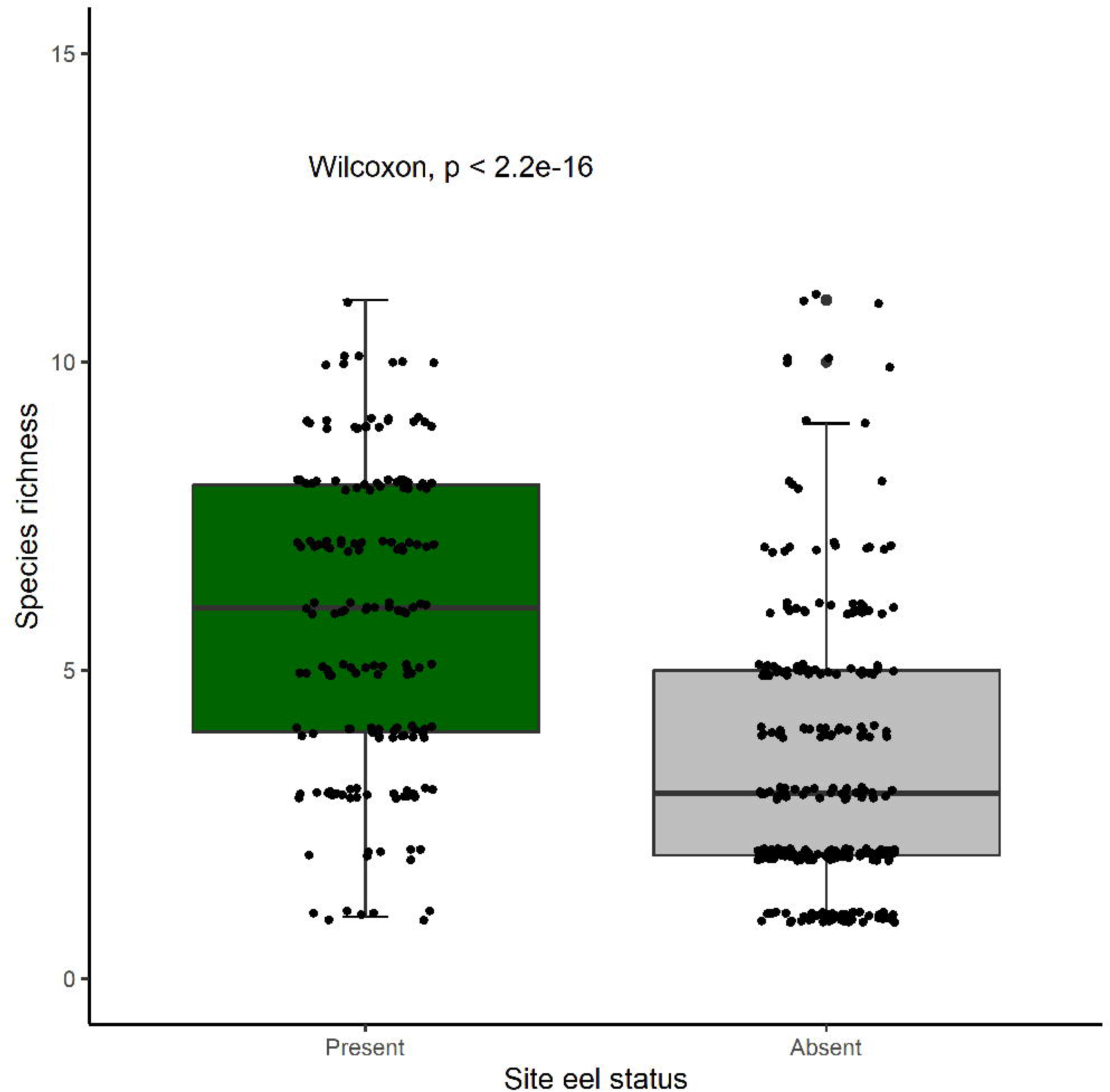
Boxplot showing the overall species richness (excluding eel) by sample at sites classified as eel present (Green) and eel absent (Grey), Wilcoxon rank sum test significance value is embedded above.

**Figure 6.**
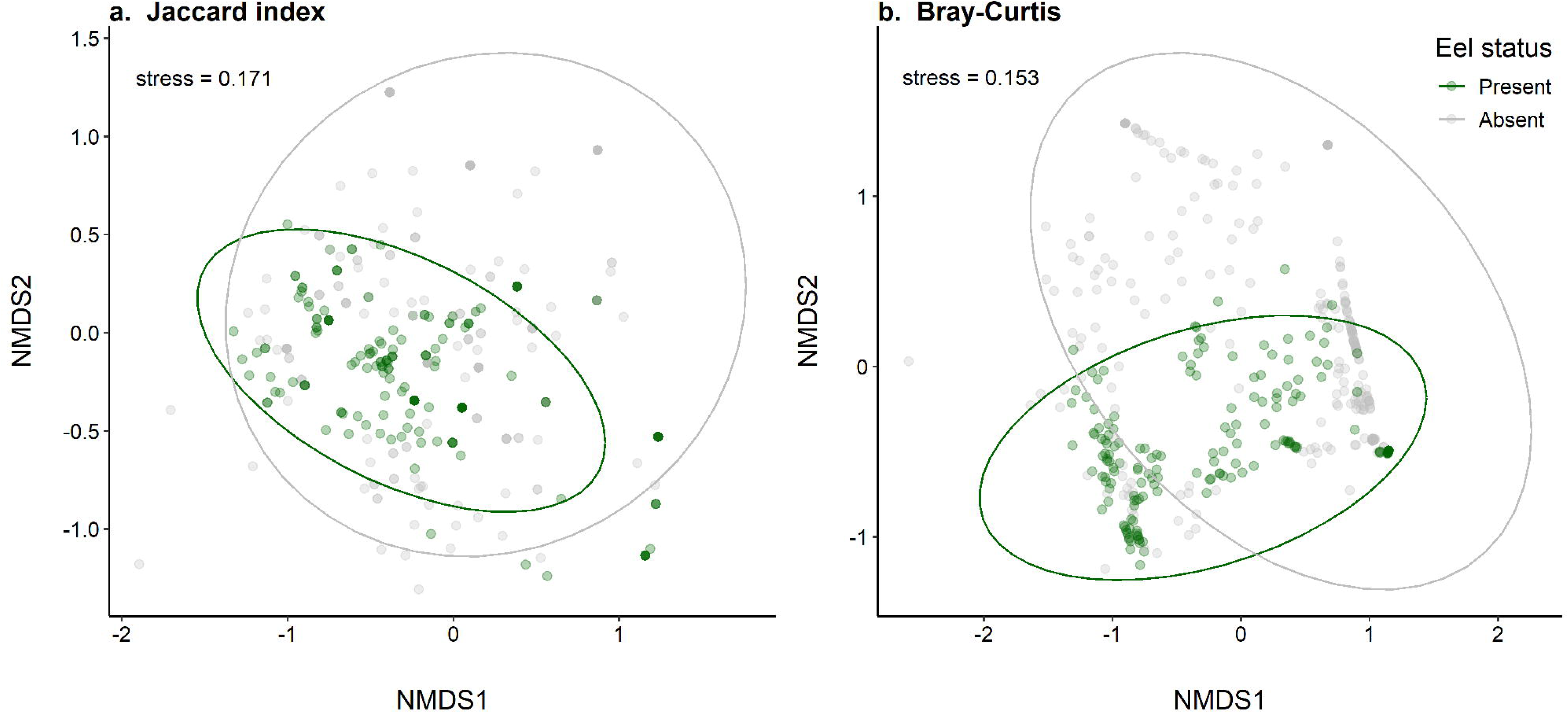
NMDS visualising the differences in fish community (excluding eel) between sites designated as eel present (green) and absent (grey). Metrics are based on measures of beta diversity including Jaccard index (a.) and Bray-Curtis (b.) including 95% confidence interval ellipses.

## 4. Discussion

### 4.1 Confidence in absence for decision-making

We developed and applied the CIADM framework (Figure 1) to eDNA metabarcoding data to provide species-specific ‘confidence in absence’ estimations given non-detection (Figure 4). The input data *n* (sample / biological replicates) and *m* (PCR / technical replicates) were based on real-world empirical data from 44 pumping stations (Figure 2), and the assumption that we over sampled was met (i.e., multiple >99% values on both axis of CIADM output matrix). Indeed, with three PCR replicates, ≥6 samples would have been sufficient to obtain the highest category of confidence (>99%) assigned in the model for our target species *A. anguilla*. However, with increased PCR replication (≥7) the same confidence could have been attained with only three sample replicates (Figure 4). Other studies have highlighted the importance of both sample and PCR replication in reducing the risk of false negatives (Ficetola *et al*., 2015; Fukaya *et al*., 2022), with optimal strategies noted to vary between river morphologies (Bylemans *et al*., 2018). In this case study, the CIADM framework considered the levels of replication required to apply evidence-based assessment of eel presence and absence (with confidence levels) to inform prioritisation at water pumping stations. We identified 27 eel absent sites (with >99% level of confidence), which means limited resources can be prioritised on remediating safe downstream passage, as per EC eel regulations (1100/2007), at the 17 eel positive sites. In highly variable and understudied catchments with rare and elusive priority species, this is key to ensuring policy targets are effectively met.

When compared to traditional methods, eDNA metabarcoding has high sensitivity in pumped (Griffiths *et al*., 2020) and heavily managed (McDevitt *et al*., 2019) catchments. However, this still does not address the issue of uncertainty with regard to false negatives and using eDNA based outputs for decision-making (Jerde, 2019). It has been suggested that current limitations of eDNA monitoring are not barriers to implementation, the issue is that we lack the tools to account for inherent uncertainty (Sepulveda *et al*., 2020). This is reflected in pumped river catchments, where knowledge of European eel presence is urgently needed to inform targeted conservation efforts. Our case study addresses this by providing a framework to aid decision-making based on priority species presence and confidence in absence, however as researchers, we can only carry out phase 1 and 2 of CIADM (Figure 4). Phase 3 requires the designation of *x* (required confidence for decision-making) by stakeholders/decision makers. If for example, x = >99%, then the 139 combinations of resampling strategies to sustain that level of confidence would apply (Figure 4). The acceptable value of *x* however will vary, depending on specific policy criteria or stakeholder risk tolerance. We recommend that CIADM should be applied as an iterative framework, and resampling data fed back into the model to ensure dynamic confidence values over time. This will ensure that if adopted long-term, the framework remains robust to environmental (i.e. flow, climatic) and protocol (i.e. lab/field workflow) changes. By developing an adaptive approach which involves stakeholder input, the CIADM framework fulfils criteria for successful integration into aquatic monitoring (Blancher *et al*., 2022).

### 4.2 Optimising single-site replication

In addition to our case-study providing a range of strategies to attain estimated confidence levels (Figure 4), it highlights the relative importance of single-site biological (sample) and technical (PCR) replication more generally. While increasing PCR replication is considered an effective way to increase detectability (Ficetola *et al*., 2015; Shirazi *et al*., 2021), our data highlight the importance of single-site sample replication to ensure a high confidence in absence can be attained. Indeed, with enough initial samples, lab effort can be adjusted to obtain the required *x* value for decision-making (Figure 4, Supplementary Information). As an example, we would recommend taking a minimum of three sample replicates if resampling the 44 sites used in this case study, to ensure that a desired level of confidence (*x*) can be reached with sufficient technical (PCR) replication. This is important, as it highlights that biological replication is still required in non-spatially replicated survey designs in order to rule out false negatives, i.e. simply failing to collect DNA for a species that is present. It must be noted however, that the optimal level of biological (sample) and technical (PCR) replication will likely depend on cost-benefit analysis (Fukaya *et al*., 2022), but that was beyond the scope of this study, given it is largely dependent on study specific logistics.

It should be noted that while increasing PCR replication improves the probability of detection, as observed in our case-study, in metabarcoding studies detectability is also influenced by sample read depth (Shirazi *et al*., 2021). Many eDNA metabarcoding studies pool PCR replicates as one multiplex (Blabolil *et al*., 2022; Griffiths *et al*., 2020; Hänfling *et al*., 2016). However, relative sample read depth should be considered when pooling many PCR replicates, as significant reductions in read depth may hamper the increased detectability gained from adding additional replicates. In our study, three PCR replicates were pooled and ran for each sample, as is standard with this 12S workflow (Blabolil *et al*., 2022; Di Muri *et al*., 2020; Griffiths *et al*., 2020; Hänfling *et al*., 2016). While this means that individually sequenced PCR replicates had comparatively increased read depth, this is not a concern, since the primary application of these were to confirm the conditional probabilities of presence in sample replicates (p*_s_*) and detection in PCR replicates (p*_r_*). The number of assigned sequence reads therefore do not influence CIADM outputs, although non-detection due to inadequate read depth would. Therefore, if a future strategy to increase PCR replication to enable fewer samples is preferred, we recommend that relative sample read depth should be adjusted for.

If false positive rates are found to be high (i.e. contamination is identified), increased levels of replication can lead to high numbers of false positive detections when left unaccounted for (Ficetola *et al*., 2015). Our framework does not inherently account for false positives, and study designs which assume no false positive results can overestimate species occupancy (Buxton *et al*., 2017). One option to reduce the risk of false positives is to implement a conservative occupancy approach alongside CIADM, such as discarding species only detected in single replicates at a site. However, many true positives can also be discarded with this approach (Ficetola *et al*., 2015), especially for rare and elusive species with an inherently low detectability (i.e. p*_s_* and p*_r_*), so there is a trade-off here. We therefore suggest that under normal circumstances CIADM is implemented alongside stringent control measures (i.e. Lab and Field blanks), in addition to appropriate data filtering, including low-frequency read thresholds (Hänfling *et al*., 2016). Designations with target species which occur in blanks, or detections which fall below thresholds, can then be regarded as ‘uncertain’ and resampled with an informed strategy. Additionally, by working closely with stakeholders, situation specific decision-support schematics can also account for uncertain classifications based on specific management plans (Sepulveda *et al*., 2020).

### 4.3 Wider applications

Confident assignments of site status can also support further downstream analysis. In this case study, eel positive sites had significantly higher species richness in fish communities than eel absent sites (Figure 5). Since pumping stations can lead to a reduction in connectivity (Kroes *et al*., 2020), it may be that increased species richness is indicative of better connected systems, and therefore more accessible for eels and other species. Alternatively, habitat quality may be better, and therefore able to support an improved fish community (including eels) more generally. Regardless of the reason, eel priority sites were more species rich, and thus conservation measures targeted at eels, such as measures to prevent entrainment in hazardous pumps (Buysse *et al*., 2015), will also benefit overall fish diversity (Itakura *et al*., 2020). The overall species community also varied significantly between eel and non-eel sites (Figure 6), therefore further research could investigate species co-occurrence and associations which could in turn further support prioritisation criteria.

The main focus of this case-study was to assign *A. anguilla* status, and as a result, the PCR replication tests to determine p*_r_* were carried out at a specifically selected eel positive site with moderate standard detection (50% of samples initially positive). While six other species were present at this site, their frequency of detection was not prioritised during site selection. This dataset is therefore biased towards our target species, but CIADM was extended to the additional six species during this study, producing a similar range of strategies to attain high levels of confidence (Supplementary information). Going forward, the CIADM framework could be similarly extended to other target species at a variety of sites/structures as required. However, the initial input data in CIADM^1^ should be representative of study systems and target species. In addition, future iterations of this framework could be built upon by factoring in additional hydromorphological variables. In the current study, conditional probabilities of detection were constant. However, by considering habitat covariates future iterations of this model may be more robust across a wider range of applications.

## 5. Conclusions

We developed a framework (CIADM) which can assign ‘confidence in absence’ given non-detection of species from eDNA metabarcoding data. In addition to enabling increased confidence in decision-making, this framework can be used to optimise resampling strategies thereby avoiding unnecessary oversampling and wasted resources. Furthermore, this framework is based on real-world empirical data and is iterative in its application – allowing for an adaptive approach to implementation and stakeholder input. The present case study successfully applied CIADM to single visit data to ascertain presence and confidence in absence for seven species, including critically endangered *A. anguilla*, at pumping stations, and produced a range of appropriate resampling strategies (139) given *x* (required confidence for decision-making). Findings suggest that, in this instance, single-site biological replication is essential to achieve high levels of confidence in absence, however PCR replication can also drive confidence once a minimum sampling threshold is reached. In our case, three single point water samples would be the minimum to ensure >99% confidence is attainable (with ≥7 PCR replicates) for *A. anguilla*. We propose this novel framework could be applied and extended to inform management decisions and survey designs relating to a range of priority species and structures.

## DATA AVAILABILITY STATEMENT

All scripts and corresponding data have been archived and made available at Zenodo: https://doi.org/10.5281/zenodo.11182844

