## Supplementary information for "Proving a negative; estimating species ‘Confidence in Absence for Decision-Making’ (CIADM) using environmental DNA monitoring"

***Anguilla anguilla***

|  | **S1** | **S2** | **S3** | **S4** | **S5** | **S6** | **S7** | **S8** | **S9** | **S10** |
| --- | --- | --- | --- | --- | --- | --- | --- | --- | --- | --- |
| **R1** | 0.67 | 0.72 | 0.77 | 0.81 | 0.85 | 0.88 | 0.90 | 0.92 | 0.94 | 0.95 |
| **R2** | 0.72 | 0.81 | 0.87 | 0.92 | 0.95 | 0.97 | 0.98 | 0.99 | >0.99 | >0.99 |
| **R3** | 0.76 | 0.87 | 0.93 | 0.97 | 0.98 | >0.99 | >0.99 | >0.99 | >0.99 | >0.99 |
| **R4** | 0.80 | 0.91 | 0.96 | 0.98 | >0.99 | >0.99 | >0.99 | >0.99 | >0.99 | >0.99 |
| **R5** | 0.83 | 0.94 | 0.98 | >0.99 | >0.99 | >0.99 | >0.99 | >0.99 | >0.99 | >0.99 |
| **R6** | 0.85 | 0.95 | 0.99 | >0.99 | >0.99 | >0.99 | >0.99 | >0.99 | >0.99 | >0.99 |
| **R7** | 0.87 | 0.96 | >0.99 | >0.99 | >0.99 | >0.99 | >0.99 | >0.99 | >0.99 | >0.99 |
| **R8** | 0.88 | 0.97 | >0.99 | >0.99 | >0.99 | >0.99 | >0.99 | >0.99 | >0.99 | >0.99 |
| **R9** | 0.89 | 0.98 | >0.99 | >0.99 | >0.99 | >0.99 | >0.99 | >0.99 | >0.99 | >0.99 |
| **R10** | 0.90 | 0.98 | >0.99 | >0.99 | >0.99 | >0.99 | >0.99 | >0.99 | >0.99 | >0.99 |
| **R11** | 0.91 | 0.98 | >0.99 | >0.99 | >0.99 | >0.99 | >0.99 | >0.99 | >0.99 | >0.99 |
| **R12** | 0.91 | 0.98 | >0.99 | >0.99 | >0.99 | >0.99 | >0.99 | >0.99 | >0.99 | >0.99 |
| **R13** | 0.91 | 0.99 | >0.99 | >0.99 | >0.99 | >0.99 | >0.99 | >0.99 | >0.99 | >0.99 |
| **R14** | 0.92 | 0.99 | >0.99 | >0.99 | >0.99 | >0.99 | >0.99 | >0.99 | >0.99 | >0.99 |
| **R15** | 0.92 | 0.99 | >0.99 | >0.99 | >0.99 | >0.99 | >0.99 | >0.99 | >0.99 | >0.99 |
| **R16** | 0.92 | 0.99 | >0.99 | >0.99 | >0.99 | >0.99 | >0.99 | >0.99 | >0.99 | >0.99 |
| **R17** | 0.92 | 0.99 | >0.99 | >0.99 | >0.99 | >0.99 | >0.99 | >0.99 | >0.99 | >0.99 |
| **R18** | 0.92 | 0.99 | >0.99 | >0.99 | >0.99 | >0.99 | >0.99 | >0.99 | >0.99 | >0.99 |
| **R19** | 0.92 | 0.99 | >0.99 | >0.99 | >0.99 | >0.99 | >0.99 | >0.99 | >0.99 | >0.99 |
| **R20** | 0.92 | 0.99 | >0.99 | >0.99 | >0.99 | >0.99 | >0.99 | >0.99 | >0.99 | >0.99 |

***Cobitis taenia***

|  | **S1** | **S2** | **S3** | **S4** | **S5** | **S6** | **S7** | **S8** | **S9** | **S10** |
| --- | --- | --- | --- | --- | --- | --- | --- | --- | --- | --- |
| **R1** | 0.80 | 0.82 | 0.85 | 0.87 | 0.89 | 0.90 | 0.92 | 0.93 | 0.94 | 0.95 |
| **R2** | 0.82 | 0.87 | 0.90 | 0.93 | 0.95 | 0.96 | 0.97 | 0.98 | 0.99 | 0.99 |
| **R3** | 0.85 | 0.90 | 0.94 | 0.96 | 0.98 | 0.99 | >0.99 | >0.99 | >0.99 | >0.99 |
| **R4** | 0.87 | 0.93 | 0.96 | 0.98 | 0.99 | >0.99 | >0.99 | >0.99 | >0.99 | >0.99 |
| **R5** | 0.89 | 0.95 | 0.98 | 0.99 | >0.99 | >0.99 | >0.99 | >0.99 | >0.99 | >0.99 |
| **R6** | 0.90 | 0.96 | 0.99 | >0.99 | >0.99 | >0.99 | >0.99 | >0.99 | >0.99 | >0.99 |
| **R7** | 0.92 | 0.97 | >0.99 | >0.99 | >0.99 | >0.99 | >0.99 | >0.99 | >0.99 | >0.99 |
| **R8** | 0.93 | 0.98 | >0.99 | >0.99 | >0.99 | >0.99 | >0.99 | >0.99 | >0.99 | >0.99 |
| **R9** | 0.94 | 0.99 | >0.99 | >0.99 | >0.99 | >0.99 | >0.99 | >0.99 | >0.99 | >0.99 |
| **R10** | 0.95 | 0.99 | >0.99 | >0.99 | >0.99 | >0.99 | >0.99 | >0.99 | >0.99 | >0.99 |
| **R11** | 0.95 | >0.99 | >0.99 | >0.99 | >0.99 | >0.99 | >0.99 | >0.99 | >0.99 | >0.99 |
| **R12** | 0.96 | >0.99 | >0.99 | >0.99 | >0.99 | >0.99 | >0.99 | >0.99 | >0.99 | >0.99 |
| **R13** | 0.97 | >0.99 | >0.99 | >0.99 | >0.99 | >0.99 | >0.99 | >0.99 | >0.99 | >0.99 |
| **R14** | 0.97 | >0.99 | >0.99 | >0.99 | >0.99 | >0.99 | >0.99 | >0.99 | >0.99 | >0.99 |
| **R15** | 0.98 | >0.99 | >0.99 | >0.99 | >0.99 | >0.99 | >0.99 | >0.99 | >0.99 | >0.99 |
| **R16** | 0.98 | >0.99 | >0.99 | >0.99 | >0.99 | >0.99 | >0.99 | >0.99 | >0.99 | >0.99 |
| **R17** | 0.98 | >0.99 | >0.99 | >0.99 | >0.99 | >0.99 | >0.99 | >0.99 | >0.99 | >0.99 |
| **R18** | 0.99 | >0.99 | >0.99 | >0.99 | >0.99 | >0.99 | >0.99 | >0.99 | >0.99 | >0.99 |
| **R19** | 0.99 | >0.99 | >0.99 | >0.99 | >0.99 | >0.99 | >0.99 | >0.99 | >0.99 | >0.99 |
| **R20** | 0.99 | >0.99 | >0.99 | >0.99 | >0.99 | >0.99 | >0.99 | >0.99 | >0.99 | >0.99 |

***Rutilus rutilus***

|  | **S1** | **S2** | **S3** | **S4** | **S5** | **S6** | **S7** | **S8** | **S9** | **S10** |
| --- | --- | --- | --- | --- | --- | --- | --- | --- | --- | --- |
| **R1** | 0.50 | 0.68 | 0.82 | 0.91 | 0.95 | 0.98 | 0.99 | >0.99 | >0.99 | >0.99 |
| **R2** | 0.63 | 0.86 | 0.96 | 0.99 | >0.99 | >0.99 | >0.99 | >0.99 | >0.99 | >0.99 |
| **R3** | 0.70 | 0.92 | 0.98 | >0.99 | >0.99 | >0.99 | >0.99 | >0.99 | >0.99 | >0.99 |
| **R4** | 0.73 | 0.94 | 0.99 | >0.99 | >0.99 | >0.99 | >0.99 | >0.99 | >0.99 | >0.99 |
| **R5** | 0.75 | 0.95 | >0.99 | >0.99 | >0.99 | >0.99 | >0.99 | >0.99 | >0.99 | >0.99 |
| **R6** | 0.75 | 0.95 | >0.99 | >0.99 | >0.99 | >0.99 | >0.99 | >0.99 | >0.99 | >0.99 |
| **R7** | 0.75 | 0.95 | >0.99 | >0.99 | >0.99 | >0.99 | >0.99 | >0.99 | >0.99 | >0.99 |
| **R8** | 0.75 | 0.95 | >0.99 | >0.99 | >0.99 | >0.99 | >0.99 | >0.99 | >0.99 | >0.99 |
| **R9** | 0.75 | 0.95 | >0.99 | >0.99 | >0.99 | >0.99 | >0.99 | >0.99 | >0.99 | >0.99 |
| **R10** | 0.75 | 0.95 | >0.99 | >0.99 | >0.99 | >0.99 | >0.99 | >0.99 | >0.99 | >0.99 |
| **R11** | 0.75 | 0.95 | >0.99 | >0.99 | >0.99 | >0.99 | >0.99 | >0.99 | >0.99 | >0.99 |
| **R12** | 0.75 | 0.95 | >0.99 | >0.99 | >0.99 | >0.99 | >0.99 | >0.99 | >0.99 | >0.99 |
| **R13** | 0.75 | 0.95 | >0.99 | >0.99 | >0.99 | >0.99 | >0.99 | >0.99 | >0.99 | >0.99 |
| **R14** | 0.75 | 0.95 | >0.99 | >0.99 | >0.99 | >0.99 | >0.99 | >0.99 | >0.99 | >0.99 |
| **R15** | 0.75 | 0.95 | >0.99 | >0.99 | >0.99 | >0.99 | >0.99 | >0.99 | >0.99 | >0.99 |
| **R16** | 0.75 | 0.95 | >0.99 | >0.99 | >0.99 | >0.99 | >0.99 | >0.99 | >0.99 | >0.99 |
| **R17** | 0.75 | 0.95 | >0.99 | >0.99 | >0.99 | >0.99 | >0.99 | >0.99 | >0.99 | >0.99 |
| **R18** | 0.75 | 0.95 | >0.99 | >0.99 | >0.99 | >0.99 | >0.99 | >0.99 | >0.99 | >0.99 |
| **R19** | 0.75 | 0.95 | >0.99 | >0.99 | >0.99 | >0.99 | >0.99 | >0.99 | >0.99 | >0.99 |
| **R20** | 0.75 | 0.95 | >0.99 | >0.99 | >0.99 | >0.99 | >0.99 | >0.99 | >0.99 | >0.99 |

***Scardinius erythrophthalmus***

|  | **S1** | **S2** | **S3** | **S4** | **S5** | **S6** | **S7** | **S8** | **S9** | **S10** |
| --- | --- | --- | --- | --- | --- | --- | --- | --- | --- | --- |
| **R1** | 0.87 | 0.95 | 0.98 | >0.99 | >0.99 | >0.99 | >0.99 | >0.99 | >0.99 | >0.99 |
| **R2** | 0.90 | 0.97 | >0.99 | >0.99 | >0.99 | >0.99 | >0.99 | >0.99 | >0.99 | >0.99 |
| **R3** | 0.90 | 0.97 | >0.99 | >0.99 | >0.99 | >0.99 | >0.99 | >0.99 | >0.99 | >0.99 |
| **R4** | 0.90 | 0.97 | >0.99 | >0.99 | >0.99 | >0.99 | >0.99 | >0.99 | >0.99 | >0.99 |
| **R5** | 0.90 | 0.97 | >0.99 | >0.99 | >0.99 | >0.99 | >0.99 | >0.99 | >0.99 | >0.99 |
| **R6** | 0.90 | 0.97 | >0.99 | >0.99 | >0.99 | >0.99 | >0.99 | >0.99 | >0.99 | >0.99 |
| **R7** | 0.90 | 0.97 | >0.99 | >0.99 | >0.99 | >0.99 | >0.99 | >0.99 | >0.99 | >0.99 |
| **R8** | 0.90 | 0.97 | >0.99 | >0.99 | >0.99 | >0.99 | >0.99 | >0.99 | >0.99 | >0.99 |
| **R9** | 0.90 | 0.97 | >0.99 | >0.99 | >0.99 | >0.99 | >0.99 | >0.99 | >0.99 | >0.99 |
| **R10** | 0.90 | 0.97 | >0.99 | >0.99 | >0.99 | >0.99 | >0.99 | >0.99 | >0.99 | >0.99 |
| **R11** | 0.90 | 0.97 | >0.99 | >0.99 | >0.99 | >0.99 | >0.99 | >0.99 | >0.99 | >0.99 |
| **R12** | 0.90 | 0.97 | >0.99 | >0.99 | >0.99 | >0.99 | >0.99 | >0.99 | >0.99 | >0.99 |
| **R13** | 0.90 | 0.97 | >0.99 | >0.99 | >0.99 | >0.99 | >0.99 | >0.99 | >0.99 | >0.99 |
| **R14** | 0.90 | 0.97 | >0.99 | >0.99 | >0.99 | >0.99 | >0.99 | >0.99 | >0.99 | >0.99 |
| **R15** | 0.90 | 0.97 | >0.99 | >0.99 | >0.99 | >0.99 | >0.99 | >0.99 | >0.99 | >0.99 |
| **R16** | 0.90 | 0.97 | >0.99 | >0.99 | >0.99 | >0.99 | >0.99 | >0.99 | >0.99 | >0.99 |
| **R17** | 0.90 | 0.97 | >0.99 | >0.99 | >0.99 | >0.99 | >0.99 | >0.99 | >0.99 | >0.99 |
| **R18** | 0.90 | 0.97 | >0.99 | >0.99 | >0.99 | >0.99 | >0.99 | >0.99 | >0.99 | >0.99 |
| **R19** | 0.90 | 0.97 | >0.99 | >0.99 | >0.99 | >0.99 | >0.99 | >0.99 | >0.99 | >0.99 |
| **R20** | 0.90 | 0.97 | >0.99 | >0.99 | >0.99 | >0.99 | >0.99 | >0.99 | >0.99 | >0.99 |

***Esox lucius***

|  | **S1** | **S2** | **S3** | **S4** | **S5** | **S6** | **S7** | **S8** | **S9** | **S10** |
| --- | --- | --- | --- | --- | --- | --- | --- | --- | --- | --- |
| **R1** | 0.50 | 0.71 | 0.85 | 0.93 | 0.97 | 0.99 | >0.99 | >0.99 | >0.99 | >0.99 |
| **R2** | 0.71 | 0.93 | 0.99 | >0.99 | >0.99 | >0.99 | >0.99 | >0.99 | >0.99 | >0.99 |
| **R3** | 0.85 | 0.99 | >0.99 | >0.99 | >0.99 | >0.99 | >0.99 | >0.99 | >0.99 | >0.99 |
| **R4** | 0.93 | >0.99 | >0.99 | >0.99 | >0.99 | >0.99 | >0.99 | >0.99 | >0.99 | >0.99 |
| **R5** | 0.97 | >0.99 | >0.99 | >0.99 | >0.99 | >0.99 | >0.99 | >0.99 | >0.99 | >0.99 |
| **R6** | 0.99 | >0.99 | >0.99 | >0.99 | >0.99 | >0.99 | >0.99 | >0.99 | >0.99 | >0.99 |
| **R7** | >0.99 | >0.99 | >0.99 | >0.99 | >0.99 | >0.99 | >0.99 | >0.99 | >0.99 | >0.99 |
| **R8** | >0.99 | >0.99 | >0.99 | >0.99 | >0.99 | >0.99 | >0.99 | >0.99 | >0.99 | >0.99 |
| **R9** | >0.99 | >0.99 | >0.99 | >0.99 | >0.99 | >0.99 | >0.99 | >0.99 | >0.99 | >0.99 |
| **R10** | >0.99 | >0.99 | >0.99 | >0.99 | >0.99 | >0.99 | >0.99 | >0.99 | >0.99 | >0.99 |
| **R11** | >0.99 | >0.99 | >0.99 | >0.99 | >0.99 | >0.99 | >0.99 | >0.99 | >0.99 | >0.99 |
| **R12** | >0.99 | >0.99 | >0.99 | >0.99 | >0.99 | >0.99 | >0.99 | >0.99 | >0.99 | >0.99 |
| **R13** | >0.99 | >0.99 | >0.99 | >0.99 | >0.99 | >0.99 | >0.99 | >0.99 | >0.99 | >0.99 |
| **R14** | >0.99 | >0.99 | >0.99 | >0.99 | >0.99 | >0.99 | >0.99 | >0.99 | >0.99 | >0.99 |
| **R15** | >0.99 | >0.99 | >0.99 | >0.99 | >0.99 | >0.99 | >0.99 | >0.99 | >0.99 | >0.99 |
| **R16** | >0.99 | >0.99 | >0.99 | >0.99 | >0.99 | >0.99 | >0.99 | >0.99 | >0.99 | >0.99 |
| **R17** | >0.99 | >0.99 | >0.99 | >0.99 | >0.99 | >0.99 | >0.99 | >0.99 | >0.99 | >0.99 |
| **R18** | >0.99 | >0.99 | >0.99 | >0.99 | >0.99 | >0.99 | >0.99 | >0.99 | >0.99 | >0.99 |
| **R19** | >0.99 | >0.99 | >0.99 | >0.99 | >0.99 | >0.99 | >0.99 | >0.99 | >0.99 | >0.99 |
| **R20** | >0.99 | >0.99 | >0.99 | >0.99 | >0.99 | >0.99 | >0.99 | >0.99 | >0.99 | >0.99 |

***Gasterosteus aculeatus***

|  | **S1** | **S2** | **S3** | **S4** | **S5** | **S6** | **S7** | **S8** | **S9** | **S10** |
| --- | --- | --- | --- | --- | --- | --- | --- | --- | --- | --- |
| **R1** | 0.33 | 0.46 | 0.59 | 0.71 | 0.81 | 0.88 | 0.92 | 0.95 | 0.97 | 0.98 |
| **R2** | 0.46 | 0.71 | 0.88 | 0.95 | 0.98 | >0.99 | >0.99 | >0.99 | >0.99 | >0.99 |
| **R3** | 0.59 | 0.88 | 0.97 | >0.99 | >0.99 | >0.99 | >0.99 | >0.99 | >0.99 | >0.99 |
| **R4** | 0.71 | 0.95 | >0.99 | >0.99 | >0.99 | >0.99 | >0.99 | >0.99 | >0.99 | >0.99 |
| **R5** | 0.81 | 0.98 | >0.99 | >0.99 | >0.99 | >0.99 | >0.99 | >0.99 | >0.99 | >0.99 |
| **R6** | 0.88 | >0.99 | >0.99 | >0.99 | >0.99 | >0.99 | >0.99 | >0.99 | >0.99 | >0.99 |
| **R7** | 0.92 | >0.99 | >0.99 | >0.99 | >0.99 | >0.99 | >0.99 | >0.99 | >0.99 | >0.99 |
| **R8** | 0.95 | >0.99 | >0.99 | >0.99 | >0.99 | >0.99 | >0.99 | >0.99 | >0.99 | >0.99 |
| **R9** | 0.97 | >0.99 | >0.99 | >0.99 | >0.99 | >0.99 | >0.99 | >0.99 | >0.99 | >0.99 |
| **R10** | 0.98 | >0.99 | >0.99 | >0.99 | >0.99 | >0.99 | >0.99 | >0.99 | >0.99 | >0.99 |
| **R11** | 0.99 | >0.99 | >0.99 | >0.99 | >0.99 | >0.99 | >0.99 | >0.99 | >0.99 | >0.99 |
| **R12** | >0.99 | >0.99 | >0.99 | >0.99 | >0.99 | >0.99 | >0.99 | >0.99 | >0.99 | >0.99 |
| **R13** | >0.99 | >0.99 | >0.99 | >0.99 | >0.99 | >0.99 | >0.99 | >0.99 | >0.99 | >0.99 |
| **R14** | >0.99 | >0.99 | >0.99 | >0.99 | >0.99 | >0.99 | >0.99 | >0.99 | >0.99 | >0.99 |
| **R15** | >0.99 | >0.99 | >0.99 | >0.99 | >0.99 | >0.99 | >0.99 | >0.99 | >0.99 | >0.99 |
| **R16** | >0.99 | >0.99 | >0.99 | >0.99 | >0.99 | >0.99 | >0.99 | >0.99 | >0.99 | >0.99 |
| **R17** | >0.99 | >0.99 | >0.99 | >0.99 | >0.99 | >0.99 | >0.99 | >0.99 | >0.99 | >0.99 |
| **R18** | >0.99 | >0.99 | >0.99 | >0.99 | >0.99 | >0.99 | >0.99 | >0.99 | >0.99 | >0.99 |
| **R19** | >0.99 | >0.99 | >0.99 | >0.99 | >0.99 | >0.99 | >0.99 | >0.99 | >0.99 | >0.99 |
| **R20** | >0.99 | >0.99 | >0.99 | >0.99 | >0.99 | >0.99 | >0.99 | >0.99 | >0.99 | >0.99 |

***Pungitius pungitius***

|  | **S1** | **S2** | **S3** | **S4** | **S5** | **S6** | **S7** | **S8** | **S9** | **S10** |
| --- | --- | --- | --- | --- | --- | --- | --- | --- | --- | --- |
| **R1** | 0.30 | 0.45 | 0.61 | 0.76 | 0.86 | 0.92 | 0.96 | 0.98 | 0.99 | >0.99 |
| **R2** | 0.41 | 0.69 | 0.88 | 0.96 | 0.99 | >0.99 | >0.99 | >0.99 | >0.99 | >0.99 |
| **R3** | 0.49 | 0.81 | 0.95 | 0.99 | >0.99 | >0.99 | >0.99 | >0.99 | >0.99 | >0.99 |
| **R4** | 0.54 | 0.86 | 0.97 | >0.99 | >0.99 | >0.99 | >0.99 | >0.99 | >0.99 | >0.99 |
| **R5** | 0.56 | 0.88 | 0.98 | >0.99 | >0.99 | >0.99 | >0.99 | >0.99 | >0.99 | >0.99 |
| **R6** | 0.57 | 0.89 | 0.98 | >0.99 | >0.99 | >0.99 | >0.99 | >0.99 | >0.99 | >0.99 |
| **R7** | 0.57 | 0.89 | 0.98 | >0.99 | >0.99 | >0.99 | >0.99 | >0.99 | >0.99 | >0.99 |
| **R8** | 0.57 | 0.89 | 0.98 | >0.99 | >0.99 | >0.99 | >0.99 | >0.99 | >0.99 | >0.99 |
| **R9** | 0.57 | 0.89 | 0.98 | >0.99 | >0.99 | >0.99 | >0.99 | >0.99 | >0.99 | >0.99 |
| **R10** | 0.57 | 0.89 | 0.98 | >0.99 | >0.99 | >0.99 | >0.99 | >0.99 | >0.99 | >0.99 |
| **R11** | 0.57 | 0.89 | 0.98 | >0.99 | >0.99 | >0.99 | >0.99 | >0.99 | >0.99 | >0.99 |
| **R12** | 0.57 | 0.89 | 0.98 | >0.99 | >0.99 | >0.99 | >0.99 | >0.99 | >0.99 | >0.99 |
| **R13** | 0.57 | 0.89 | 0.98 | >0.99 | >0.99 | >0.99 | >0.99 | >0.99 | >0.99 | >0.99 |
| **R14** | 0.57 | 0.89 | 0.98 | >0.99 | >0.99 | >0.99 | >0.99 | >0.99 | >0.99 | >0.99 |
| **R15** | 0.57 | 0.89 | 0.98 | >0.99 | >0.99 | >0.99 | >0.99 | >0.99 | >0.99 | >0.99 |
| **R16** | 0.57 | 0.89 | 0.98 | >0.99 | >0.99 | >0.99 | >0.99 | >0.99 | >0.99 | >0.99 |
| **R17** | 0.57 | 0.89 | 0.98 | >0.99 | >0.99 | >0.99 | >0.99 | >0.99 | >0.99 | >0.99 |
| **R18** | 0.57 | 0.89 | 0.98 | >0.99 | >0.99 | >0.99 | >0.99 | >0.99 | >0.99 | >0.99 |
| **R19** | 0.57 | 0.89 | 0.98 | >0.99 | >0.99 | >0.99 | >0.99 | >0.99 | >0.99 | >0.99 |
| **R20** | 0.57 | 0.89 | 0.98 | >0.99 | >0.99 | >0.99 | >0.99 | >0.99 | >0.99 | >0.99 |
